# Live imaging of the co-translational recruitment of XBP1 mRNA to the ER and its processing by diffuse, non-polarized IRE1α

**DOI:** 10.1101/2021.11.15.468613

**Authors:** Silvia Gómez-Puerta, Roberto Ferrero, Tobias Hochstoeger, Ivan Zubiri, Jeffrey A. Chao, Tomás Aragón, Franka Voigt

**Author notes:** Corresponding author, Dr. Tomás Aragón, office: 55, Pío XII Ave. – 31008 Pamplona, Dr. Franka Voigt, office: Maulbeerstrasse 66, CH – 4058 Basel.

## Abstract

Endoplasmic reticulum (ER) to nucleus homeostatic signalling, known as the unfolded protein response (UPR), relies on the non-canonical splicing of XBP1 mRNA. The molecular switch that initiates splicing is the oligomerization of the ER stress sensor and UPR endonuclease IRE1α. While IRE1α can form large clusters that have been proposed to function as XBP1 processing centers on the ER, the actual oligomeric state of active IRE1α complexes as well as the targeting mechanism that recruits XBP1 to IRE1α oligomers, remain unknown.

Here, we used a single molecule imaging approach to directly monitor the recruitment of individual XBP1 transcripts to the ER surface. We confirmed that stable ER association of unspliced XBP1 mRNA is established through HR2-dependent targeting and relies on active translation. In addition, we show that IRE1α-catalyzed splicing mobilizes XBP1 mRNA from the ER membrane in response to ER stress. Surprisingly, we find that XBP1 transcripts are not recruited into large IRE1α clusters, which only assemble upon overexpression of fluorescently-tagged IRE1α during ER stress. Our findings support a model where ribosome-engaged, ER-poised XBP1 mRNA is processed by functional IRE1α assemblies that are homogenously distributed throughout the ER membrane.

## Introduction

Cellular organization depends on the ability of cells to recruit mRNA and protein molecules to precise subcellular localizations. In eukaryotic cells, mRNA transcripts that encode membrane and secreted proteins are targeted to the endoplasmic reticulum (ER), which provides a specific biochemical environment to ensure proper folding and maturation of these classes of proteins. mRNA targeting is mediated through the co-translational recognition of a signal sequence by the signal-recognition particle (SRP) (Walter et al., 1981). SRP-ribosome-nascent chain complexes are recruited to the surface of the ER by the SRP receptor (Gilmore et al., 1982), which channels the nascent polypeptide into the ER lumen through interaction with the Sec61 translocon (Görlich et al., 1992).

The unfolded protein response (UPR) acts as a combination of quality control pathways that monitor the folding status of proteins within the ER lumen and adjust the capacity of the ER’s folding machinery (Walter & Ron, 2011). IRE1α (inositol-requiring enzyme 1 alpha) triggers the most conserved branch of the UPR (Cox et al., 1993; Mori et al., 1993). It is an ER membrane resident stress sensor that is activated by the accumulation of misfolded proteins in the ER lumen and signals ER stress through the non-canonical splicing of X-box binding protein 1 mRNA (XBP1, *HAC1* in yeast) (Sidrauski & Walter, 1997; Tirasophon et al., 1998; Yoshida et al., 2001).

Processing of unspliced XBP1 (XBP1u) mRNA is initiated upon oligomerization and trans-autophosphorylation of IRE1α (Ali et al., 2011), which leads to the allosteric activation of its cytosolic kinase and RNAse domains (Korennykh et al., 2009). Activated IRE1α excises a highly conserved 26 nucleotide intron from the XBP1 coding sequence (Calfon et al., 2002; Yoshida et al., 2001) and the severed exons are re-joined by the tRNA ligase RtcB (Jurkin et al., 2014; Kosmaczewski et al., 2014; Lu et al., 2014). Intron excision causes a translational frameshift in the spliced (XBP1s) transcript, which encodes a potent transcription factor that increases the folding capacity of the ER through a broad activation of stress response genes (Acosta-Alvear et al., 2007), including expression of ER-associated degradation (ERAD) factors (Brodsky, 2012). Beyond processing XBP1 mRNA, metazoan IRE1α is able to cleave a variety of mRNAs to initiate their rapid degradation in a pathway known as Regulated IRE1-Dependent Decay (RIDD) (Hollien et al., 2009; Hollien & Weissman, 2006). Even though RIDD has been found to play a key role in some pathological conditions, XBP1 splicing stands out as the main physiological output of IRE1 activation (Ishikawa et al., 2017).

To efficiently support rapid responses to ER stress, eukaryotic organisms display different strategies to ensure the timely encounter of IRE1α and its substrate mRNAs. In *S. cerevisiae*, acute ER stress triggers the rapid oligomerization of Ire1p into a discrete number of foci (Aragón et al., 2009; Kimata et al., 2007). *HAC1* mRNA, the yeast homolog of XBP1, is then recruited into these foci through a bipartite element that is located in the *HAC1* 3’UTR while translational repression is imposed by the *HAC1* intron itself (Aragón et al., 2009; Rüegsegger et al., 2001; van Anken et al., 2014). This swift targeting of *HAC1* mRNA to pre-formed Ire1p clusters is essential to allow a timely response to ER stress and to sustain yeast proteostasis (Pincus et al., 2010).

The activation of metazoan IRE1α has been proposed to follow the same principles that were defined in yeast. Under ER stress, ectopic, fluorescently-labeled IRE1α was found to cluster into large dynamic foci and the kinetics of cluster assembly and disassembly approximately correlated with XBP1 splicing rates (Li et al., 2010). Yet, there is no direct evidence that formation of large IRE1α clusters is required for splicing. Even though oligomerization of IRE1α has been proven to be the regulatory step that coordinates mRNA cleavage (Korennykh et al., 2009; Li et al., 2010) and the disruption of oligomerization interfaces has been shown to diminish RNAse activity (Karagöz et al., 2017; Sanches et al., 2014), the specific oligomeric state of splicing-competent IRE1α assemblies has not been precisely determined. In addition, only a minor fraction (~5%) of all cellular IRE1α protein concentrates in detectable foci (Belyy et al., 2019) and there is no direct evidence that they are indeed the sites of XBP1 processing at the ER.

What is more, in contrast to yeast *HAC1*, metazoan XBP1 mRNA is recruited to the ER surface through co-translational targeting that involves a peptide signal sequence and not a *cis*-acting localization element. Specifically, XBP1u transcripts encode a hydrophobic stretch (HR2) located at the C-terminal half of the protein that mimics a secretion signal (Yanagitani et al., 2009). Upon translation, this hydrophobic stretch is recognized by SRP, which delivers the nascent chain complex to the Sec61 translocon in the ER membrane (Plumb et al., 2015). Recognition of the HR2 peptide is aided by a translational pausing mechanism that has been proposed to stall the translating ribosome through high-affinity interactions with the peptide exit tunnel and thus conveys stability to the mRNA-ribosome-nascent chain complex that facilitates its delivery to the ER membrane (Kanda et al., 2016; Yanagitani et al., 2011). Such a co-translational targeting mechanism suggests that IRE1α encounters XBP1u mRNA at the Sec61 translocon, where translating ribosomes would be poised. This notion is supported by the reported interaction of IRE1α with the translocon complex as well as by crosslinking data that find IRE1α in close contact with SRP, ribosomal RNAs (rRNAs) and a subset of ER-targeted mRNAs (Acosta-Alvear et al., 2018; Plumb et al., 2015). However, this model is difficult to reconcile with a situation where IRE1α molecules are recruited into large clusters with complex topologies that are not simply 2D patches in the ER membrane but have also been described to exclude the Sec61 translocon from specific regions within the clusters (Belyy et al., 2019).

Here, we directly image the recruitment of XBP1 mRNA to IRE1α and the ER surface using a single-molecule imaging approach. We find that individual XBP1 transcripts are recruited for splicing on the ER via a translation-dependent targeting mechanism that is different from the canonical SRP-mediated recruitment and demonstrate how XBP1 mRNAs are mobilized from the ER surface upon induction of ER stress. In addition, we find that ER association depends on IRE1α cleavage activity. Using a dual-color live imaging approach, we visualize individual XBP1 mRNA transcripts together with IRE1α-GFP, which only assembles into clusters at increased expression levels and does not recruit XBP1 mRNA to these higher oligomeric assemblies. Instead, we find that lower expressed IRE1α-GFP simply outlines the ER and cleaves XBP1 mRNA in the absence of cluster formation during ER stress.

## Results

In order to directly visualize the recruitment dynamics of XBP1 mRNA, we developed a single-molecule imaging approach that takes advantage of the MS2 labeling system to detect individual reporter mRNAs in living cells (Bertrand et al., 1998). We generated a XBP1 wild-type (WT) reporter transcript that comprises the complete *M.musculus* open reading frame (ORF) as well as its complete 3’UTR (Figure 1A, red) (Calfon et al., 2002; Sugimoto et al., 2015). To enable the detection of single mRNA molecules at high signal-to-noise ratios, we further included 24 MS2 stem-loops in the 3’UTR of all reporter transcripts (Figure 1A) and made use of their specific recognition by fluorescently labeled synonymous tandem MS2 coat proteins (stdMCPs) (Bertrand et al., 1998; Wu et al., 2015).

**Figure 1.**
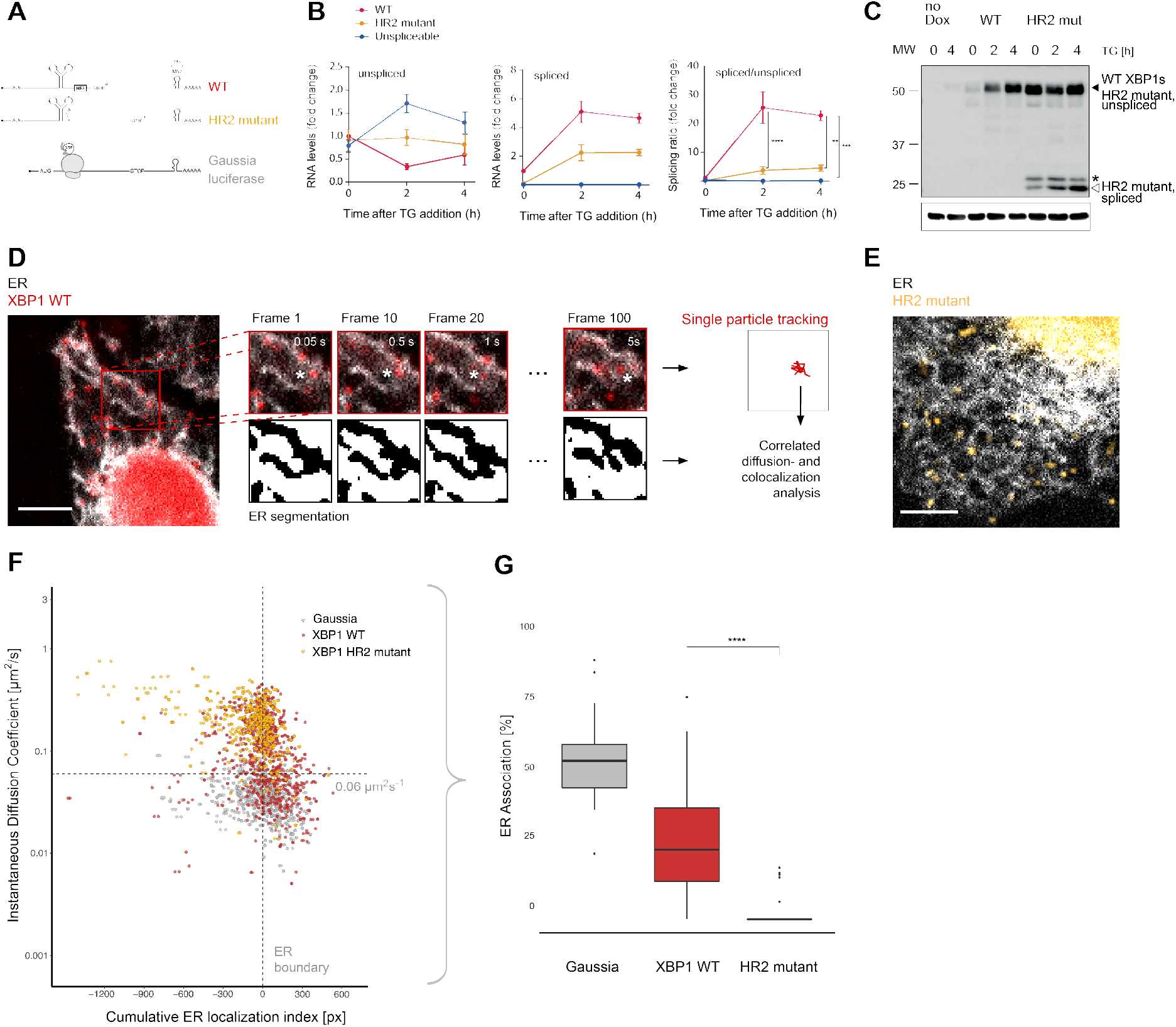
Live imaging of XBP1 mRNA recruitment to the ER. **(A)** Reporter construct design: XBP1 WT (red) features the mouse XBP1 ORF and 3’UTR and contains a 24x MS2 stem loop array for mRNA detection. XBP1 HR2 mutant (yellow) is identical to the WT construct but contains a point mutation downstream of the ER intron that renders the HR2 peptide out-offrame. The Gaussia luciferase reporter (gray) is a canonical SRP-recruited transcript and serves as positive control for ER association. **(B)** qPCR assay showing splicing of MS2-labeled XBP1 reporter transcripts upon induction of ER stress with thapsigargin (TG). HeLa cells expressing WT and HR2 mutant reporters were treated with 0.2 ug/ml doxycycline (Dox) for 15 hours before addition of 100 nM TG for indicated times. **(C)** Western blot against XBP1 protein in response to UPR activation with 100 nM TG for indicated times using an antibody that preferentially recognizes mouse over human XBP1 (human XBP1s background signal is detectable in samples w/o reporter expression = no Dox). Black triangle: 55 kDa band corresponding to endogenous and reporter WT XBP1s, which have the same size as unspliced HR2 mutant protein. White triangle: spliced HR2 mutant XBP1s protein. Asterisk (*): Short protein product present before TG treatment. (**D**) Representative live-cell image of the XBP1 WT reporter (red) in a HeLa cell expressing NLS-stdMCP-stdHalo and a fluorescent ER marker (gray). Illustration of the image analysis workflow: diffraction-limited spots (*) are individual mRNA transcripts. **(E)** Same as in (D) but expressing XBP1 HR2 mutant reporters (yellow). All scale bars = 5 μm. **(F)** Correlated diffusion and ER colocalization analysis of individual XBP1 WT (red), HR2 mutant (yellow), and Gaussia (gray) transcripts. Dots are single particles that were tracked for at least 30 frames. Y axis: Instantaneous diffusion coefficients. X axis: Cumulative ER localization index. Positive values indicate ER colocalization. **(G)** Boxplot showing ER association quantified from data shown in (F). Statistical test: unpaired t-test, p-value = 1e-8.

In parallel to the XBP1 WT reporter, we generated a frameshift mutation downstream of the ER intron (Figure 1A, yellow, HR2-mutant) that prevents synthesis of the HR2 peptide, which has been shown to be essential for non-canonical SRP-mediated translocation of XBP1u mRNA to the ER membrane (Kanda et al., 2016; Yanagitani et al., 2009, 2011). In addition, we employed a previously characterized SRP-recruited reporter (Voigt et al., 2017) to benchmark ER association of XBP1 transcripts against this established reporter construct encoding a secreted Gaussia luciferase protein (Figure 1A, gray). We next generated HeLa cell lines stably expressing these reporter transcripts under a doxycycline-inducible promoter and from single genomic loci (Weidenfeld et al., 2009). To allow detection of individual mRNA particles as diffraction limited spots in the cytoplasm of living cells, we co-expressed nuclear localization signal-encoding fluorescently labeled NLS-stdMCP-stdHalo fusion proteins (Voigt et al., 2017).

To confirm that these reporter constructs were indeed splicing competent, we first performed qPCR-based splicing assays (Figure 1B). As expected, we detected an increase in the levels of spliced XBP1 WT mRNA (red), and a transient drop in the levels of unspliced WT mRNA in response to induction of ER stress with thapsigargin (TG). Using these measurements, we calculated the splicing ratio (spliced/unspliced) as a quantitative readout of splicing efficiency. As expected, the analysis showed that the response was specific to the WT reporter and we observed almost very little increase in splicing activity for the HR2 mutant construct (yellow). In parallel, we quantified XBP1 protein levels in response to TG treatment (Figure 1C) and detected increased levels of XBP1s in cells expressing WT reporter transcripts (black triangle, Figure 1C). HR2-mutant cells produced only low levels of XBP1s in response to TG treatment (white triangle, Figure 1C) while the majority of their XBP1 protein product was still derived from unspliced HR2 mutant transcripts (black triangle, same size as WT XBP1s protein).

To assess XBP1 mRNA mobility and investigate recruitment dynamics of individual transcripts, we acquired streaming movies at fast frame rates (20 Hz) that detected XBP1 mRNAs as Halo-labeled diffraction limited spots in the cytoplasm of individual HeLa cells (Figure 1D, red). We performed single-particle tracking (SPT) over 100 consecutive frames and used the resulting particle coordinates to determine instantaneous diffusion coefficients as a measure of particle mobility (H. C. Berg, 1993; Voigt et al., 2017).

According to current models, unspliced XBP1 WT mRNA (but not the HR2 mutant) should be constitutively recruited to the ER surface for IRE1α-mediated splicing during ER stress. To investigate XBP1 mRNA association with the ER, we therefore integrated a fluorescently-labeled ER marker protein (Sec61b-SNAP) into the reporter cell lines introduced above (analogous to Belyy *et al*, 2019). We imaged dual-labeled cells using a fluorescence microscope equipped with two parallel light paths and registered cameras for simultaneous detection of mRNA and ER signal in independent channels (Supplementary Movie 1).

Next, we quantified the mobility of individual particles with respect to their ER localization and therefore assessed when an mRNA particle associates with the ER. To this aim, we segmented the ER signal and used it to generate distance maps that allowed us to correlate particle coordinates with ER boundaries (Figure 1D, right panels). In these distance maps, positions on the ER were given positive values and positions away from the ER were defined as negative. Based on particle trajectories, we determined the localization of individual transcripts throughout the entire image series and calculated cumulative ER localization indices that highlight robust localization phenotypes (Voigt et al., 2017). We combined the diffusion and ER colocalization analysis and employed it to benchmark the mobility and ER association of the XBP1 WT and HR2 mutant reporters (Figure 1E) based on the behavior of a Gaussia luciferase reporter transcript that encodes a secreted protein product and that we have previously shown to be predominantly localized to the ER (Voigt et al., 2017).

The combined analysis shows that a large fraction of XBP1 WT transcripts (Figure 1F, red dots) behaves similar to the secreted Gaussia mRNAs (Figure 1F, gray dots). Many XBP1 WT transcripts exhibit low mobility and colocalize with the ER. However, there is another population of WT reporter tracks not observed for the Gaussia reporter that it is more mobile and tends to not localize to the ER. Interestingly, the behavior of this population is exactly matched by the XBP1 HR2 mutant tracks (Figure 1F, yellow dots). These reporter mRNAs appear to have lost their ability to be recruited to the ER surface and exhibit a generally higher degree of mobility that is also apparent upon visual inspection (Figure 1F, Supplementary Movie 2). We employed the correlated diffusion and ER colocalization analysis to quantify the fraction of ER-associated particles per cell. To this aim, we used the clearly ER-associated Gaussia cluster to define cut-offs (D < 0.06 μm^2^s^-1^ and positive ER localization index, dashed lines in Figure 1F) for identification of XBP1 mRNA particles that showed a similar behavior. Based on these parameters, we found an (per cell) average of 27.4 ± 19.4 % (Mean ± SD) of all XBP1 WT and 3.1 ± 6.2 % of all HR2 mutant transcripts to be associated with the ER (Figure 1G).

To corroborate the findings from the single-particle imaging approach through an independent method, we performed flotation assays that allow separation of membrane from cytosolic fractions (Supplementary Figure 1A) (Mechler & Rabbitts, 1981). As expected, we found that XBP1 WT reporter mRNAs associated with the membrane fractions to a similar extent as the endogenous XBP1u mRNA. XBP1 HR2 mutant mRNA on the contrary, lacked membrane association and behaved like endogenous spliced XBP1s (Supplementary Figure 1B). Upon reconstitution of the original open reading frame through integration of two additional nucleotides that restore the HR2 reading frame but not the upstream part of the ORF, membrane association was restored (Supplementary Figure 1C,D). Recruitment of XBP1 reporter transcripts to the ER is therefore unambiguously linked to the expression of the HR2 peptide.

Together, these findings indicate that XBP1 WT reporter mRNA is recruited to the ER surface albeit to a lesser extent than canonical secretion-signal encoding Gaussia transcripts. Recruitment depends on the expression of the HR2 peptide since a reporter mRNA that cannot produce HR2 failed to associate with the ER. Our results are consistent with the non-canonical mechanism of XBP1 delivery to the ER and confirm that HR2 expression conveys stable ER association in a co-translational manner.

To test if translation-dependent recruitment of XBP1 transcripts to the ER membrane is indeed necessary to enable mRNA splicing, we generated an XBP1 translation site reporter that would allow us to directly monitor XBP1u translation on the ER (Figure 2A). Specifically, we used a nascent polypeptide imaging approach that relies on the expression of a well-folded protein scaffold (spaghetti monster, SM) that contains nine GCN4 antigen repeats (Eichenberger et al., in preparation; Morisaki et al., 2016; Yan et al., 2016). To quantify protein synthesis of XBP1u transcripts on the ER, we generated a XBP1u translation reporter construct that contains the GCN4-SM downstream of the UPR intron but in frame with the XBP1u ORF. Upon splicing and excision of the intron by IRE1α, the ORF changes to XBP1s and the GCN4-SM is no longer in frame. Thus, the translation site signal can only be detected prior to mRNA splicing.

**Figure 2.**
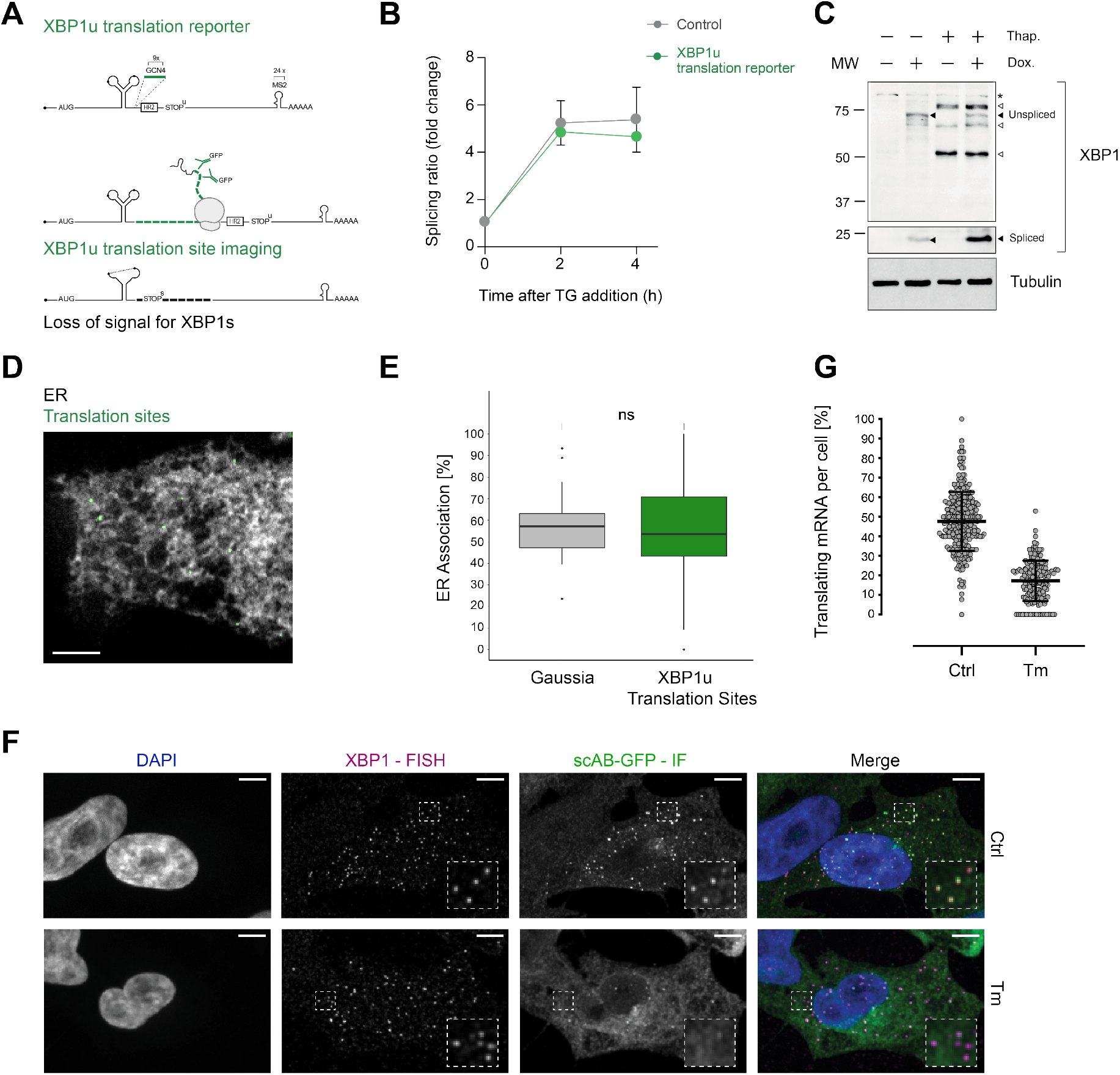
Association of XBP1u mRNA with the ER is translation dependent. **(A)** Reporter construct design and illustration of the method: XBP1u translation reporters feature a 9x GCN4 array (green) inserted into the ORF downstream of the ER intron and in frame with the XBP1u protein. Upon translation of GCN4-XBP1u, emerging GCN4 peptide repeats are recognized by GFP-labeled single-chain antibodies (scAB-GFP), which allow detection of translating ribosomes together with mRNA transcripts. Upon splicing, the reading frame is changed and GCN4 expression is lost. **(B)** qPCR-based splicing assay to test functionality of XBP1u translation reporter (green) as compared to a non GCN4-tagged control (gray). Shown is the splicing ratio (XBP1s/XBP1u) in response to induction of ER stress with 100 nM TG. **(C)** Western blot against XBP1 proteins. Spliced XBP1 appearance is dependent on reporter expression (Dox) and induction of ER stress with 100 nM TG. Black arrows: XBP1 protein products expressed upon TG and Dox treatment. White arrows: Unspecific bands present irrespective of reporter expression (Dox) in response to TG. Asterisk: Unspecific band present in all samples. **(D)** Representative live-cell image of XBP1u translation sites (green diffraction limited spots) in a HeLa cell expressing scAB-GFP and a fluorescent ER marker (gray). **(E)** Boxplot showing ER association of XBP1u translation sites (green) as compared to secreted protein encoding Gaussia mRNAs (gray) that serve as an ER-associated positive control. Statistical test: unpaired t-test, p-value = 0.49. **(F)** Combined smFISH and IF analysis for co-localization of XBP1 mRNA (magenta) and translation site signal (green) in fixed HeLa cells (DAPI = blue). The majority of translation site spots disappear upon induction of ER stress with 5 μg/ml TM for 2h. **(G)** Quantification of data shown in (F). Individual dots represent per-cell averages. Black bars show mean ± SD. All scale bars = 5 μm.

Quantitative RT-PCR as well as Western blot analysis confirmed that this construct was able to undergo splicing upon induction of ER stress (Figure 2B,C). To characterize the translational status of XBP1 mRNA in live imaging experiments, we employed GFP-fused single-chain antibodies (scAB-GFP) (Voigt et al., 2017; Yan et al., 2016) that specifically recognize GCN4 peptides and allow detection of individual translation sites as diffraction-limited spots in the cytoplasm of HeLa cells coexpressing the Sec61b-SNAP ER marker (Figure 2D, Supplementary Figure 2A, Supplementary Movie 3). For further characterization of this experimental set-up, we performed a similar dual-color live imaging experiment but this time focused on the simultaneous detection of mRNA and translation site signals. We tested whether the bright GFP signal was indeed corresponding to individual translation sites and, to this aim, first acquired the NLS-stdMCP-stdHalo mRNA in parallel with the scAB-GFP translation site signal (Supplementary Figure 2B) and then treated the cells with puromycin (PUR) to inhibit translation (Supplementary Figure 2C). Upon PUR treatment, all scAB-GFP spots disappeared, which led us to conclude that they were indeed translation sites.

We proceeded to quantify the degree of ER association observed for XBP1u translation sites in individual cells through the correlated diffusion and ER colocalization analysis introduced above (Figure 2E, Supplementary Figure 2A). Interestingly, this analysis revealed that the majority of XBP1u translation sites colocalizes with the ER (53.8 ± 22.1 %, mean per cell ± SD) and exhibits a low mobility that is comparable to the behavior of predominantly ER-localized Gaussia transcripts (Mean ER association = 57.3 ± 16.8 %) but very different from the average degree of ER association assumed by XBP1 WT transcripts (Mean ER association = 27.4 ± 19.4 %). Thus, we conclude that XBP1u reporters are indeed recruited to the ER surface in a translation-dependent manner.

As the translational frameshift induced by IRE1α-mediated splicing should abolish translation of the GCN4 repeats, we assessed the fraction of translating XBP1u transcripts in response to induction of ER stress. Accordingly, we treated the cells with tunicamycin (TM) and then quantified the degree of co-localization for XBP1u mRNA and translation site spots. To maximize detection efficiency and more accurately estimate particle numbers per cell, we performed a combined single-molecule fluorescence in-situ hybridization (smFISH) and immunofluorescence (IF) experiment in fixed cells (Figure 2F). Specifically, we used smFISH probes against the 5’-end of the *M.musculus* XBP1 ORF and co-localized their signal with the IF staining of anti-GFP antibodies that allowed detection of the scAB-GFP labeled nascent polypeptides. As expected, we found that the number of translating XBP1u particles is drastically reduced upon induction of ER stress. In fact, the majority of XBP1u translation sites disappears after only 2h TM stress and the fraction of translating mRNAs is significantly reduced from 0.47 ± 0.14 (Mean ± SD) to 0.17 ± 0.10 (Figure 2G). Taken together, these results demonstrate that the localization of XBP1 mRNA to the ER is translation-dependent and leads to splicing of XBP1u transcripts upon induction of ER stress.

Next, we investigated how ER stress affects the association of XBP1 mRNA with the ER and set out to determine how unspliced XBP1 molecules encounter IRE1α. In order to distinguish between the behavior of unspliced and spliced mRNA transcripts, we generated a reporter variant with point mutations in the 5’ and 3’ splice site of the UPR intron that maintain its stem-loop structure but render the substrate cleavage incompetent (unspliceable, dark blue, Figure 3A, Figure Supplementary Figure 3A,B) (Calfon et al., 2002; Gonzalez et al., 1999). In addition we also generated a variant lacking the intron that expresses the XBP1s protein constitutively (spliced, light blue, Figure 3A, Figure Supplementary Figure 3A,B). We performed dual-color live imaging experiments (Figure 3B, Supplementary Movie 4 and 5) and quantified reporter mobility and their degree of colocalization with the ER through correlated diffusion and ER colocalization analysis under non stress conditions.

**Figure 3.**
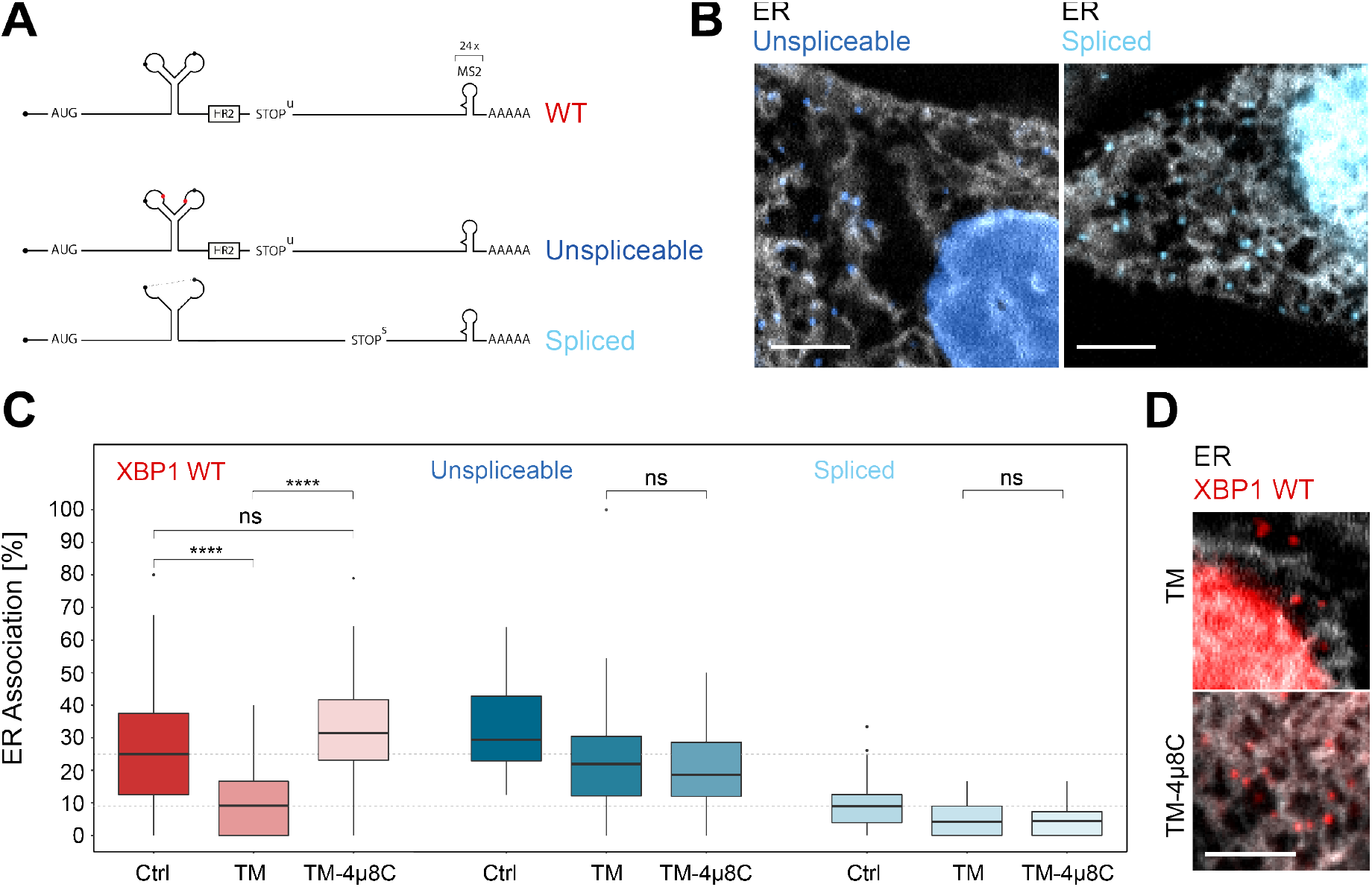
IRE1α-dependent processing and ER association of XBP1u transcripts during stress. **(A)** Reporter construct design: Unspliceable (dark blue) and spliced (light blue) reporter transcripts are identical to XBP1 WT (red) except for point mutations in the intron (unsplicable) or the complete lack of it (spliced). **(B)** Representative live-cell images of XBP1 splice site mutant reporters (blue) in HeLa cells expressing NLS-stdMCP-stdHalo and Sec61b-SNAP as ER marker (gray). **(C)** Boxplot showing quantification of ER association from correlated diffusion and ER colocalization analysis for XBP1 WT (red), unspliceable (dark blue) and spliced (light blue) reporter transcripts. Different opacities represent experimental conditions: no treatment (Ctrl), ER stress induced with 3-4h of 5 μg/ml tunicamycin (TM), ER stress induced with 3-4h of 5 μg/ml tunicamycin under IRE1α inhibition with 4μ8C (TM+4μ8C). Statistical test: unpaired t-test, p-values: (p ≥ 0.05) = ns; (p < 0.0001) = ****. **(D)** Representative live-cell images of XBP1 WT reporter constructs (red) in HeLa cells expressing NLS-stdMCP-stdHalo and Sec61b-SNAP as ER marker (gray) under ER stress (5 μg/ml TM) as well as ER stress with IRE1α inhibition (5 μg/ml TM and 50 μM 4μ8C). All scale bars = 5 μm.

As expected, unspliceable reporter transcripts often exhibit a lower mobility (Supplementary Figure 3C) and associate with the ER to a degree that is comparable to WT transcripts (Figure 3C) while spliced reporter mRNAs tend to diffuse at higher mobilities (Supplementary Figure 3D) and display a low degree of ER association (Figure 3C) comparable to cytoplasmic protein encoding mRNAs (Voigt et al., 2017).

To determine the extent of ER association for WT, unspliceable and spliced reporter transcripts, we induced ER stress with TM (5 μg/ml) at least 3h before the start of the imaging session (Figure 3D, Supplementary Movie 6). In addition, and to specifically quantify the involvement of IRE1α in the splicing reaction, we performed the same imaging experiments including 50 μM 4μ8C, a small molecule inhibitor that blocks substrate access to the active site of the IRE1α RNase domain and thereby selectively inactivates XBP1 cleavage (Supplementary Movie 7) (Cross et al., 2012). As anticipated, ER stress-induced processing of WT reporters caused a strong decrease of their mean ER association from 27.4 ± 19.4 % (mean ± SD) in the untreated condition, to 10.1 ± 9.3 % under TM treatment (Figure 3C, red). This result illustrates how, upon completion of the splicing reaction, WT mRNAs are released from the ER membrane and behave like intron-free transcripts (Figure 3C, light blue) in the absence of ER stress (10.0 ± 9.1 %). ER stress-induced mobilization of spliced WT reporter transcripts was a genuine consequence of IRE1α catalysis, since addition of 4μ8C to the TM condition restored ER association of WT reporters back to 33.2 ± 15.6 % (Figure 3C, red). Taken together, these findings suggest that IRE1α-mediated splicing drives the release of translationally active, translocon-engaged mRNAs.

In line with this notion, unspliceable reporter transcripts (Figure 3C, dark blue) not only associate with the ER to a high level (32.4 ± 13.5 %) but also fail to show a similar reduction in ER association upon TM (23.7 ± 18.4 %) and TM+4μ8C (20.3 ± 12.6 %) treatment (Figure 3C). The same holds true for the spliced reporter construct. Since it does not encode the HR2 peptide and can therefore not be delivered to the translocon, it associates with the ER to only a limited extent (10.0 ± 9.1 %). Upon induction of ER stress, its ER association rate is further reduced and similar to the unspliceable reporter, we do not observe significant changes in ER association between ER stress conditions in the absence (4.8 ± 5.0 %) and presence (4.6 ± 4.4 %) of 4μ8C.

We noticed that, for unspliceable and spliced reporters, ER stress caused a reduction of ER association when compared to untreated conditions. This effect likely results from the general inhibition of cellular translation initiation triggered by the eIF2α kinase PERK, that promotes the UPR branch of the integrated stress response (Pakos-Zebrucka et al., 2016). It is plausible that the slightly reduced levels of ER association under ER stress conditions are due to decreased recruitment of translating mRNPs to the ER surface, affecting all mRNAs to a limited extent (Voigt et al., 2017). This effect is not observed for the XBP1 WT reporter, where conversion of unspliced molecules into spliced ones is the major driver of mobilization from the ER. In summary, these experiments demonstrate that IRE1α activity is not required for ER association of XBP1 reporter mRNAs, but that IRE1α-mediated catalysis (UPR splicing) determines the release of spliced mRNA molecules to the cytosol.

In combination, our findings support a model where IRE1α-mediated splicing is instrumental for the mobilization of XBP1 transcripts that are anchored to the ER in a translation-dependent manner. Based on this hypothesis, we sought to further investigate and visualize the sites of XBP1 processing on the ER membrane. To this aim, we developed an approach that allowed us to detect IRE1α in the reporter transcript-expressing HeLa cell lines introduced above. We knocked out the endogenous IRE1α using CRISPR/Cas9 and reconstituted its expression with a GFP-tagged IRE1α protein (Figure 4A). Based on the previously published design of a splicing competent IRE1α-GFP construct (Belyy et al., 2019), we introduced a GFP moiety in between the lumenal and kinase/RNase domains on the cytoplasmic site of the transmembrane protein.

**Figure 4.**
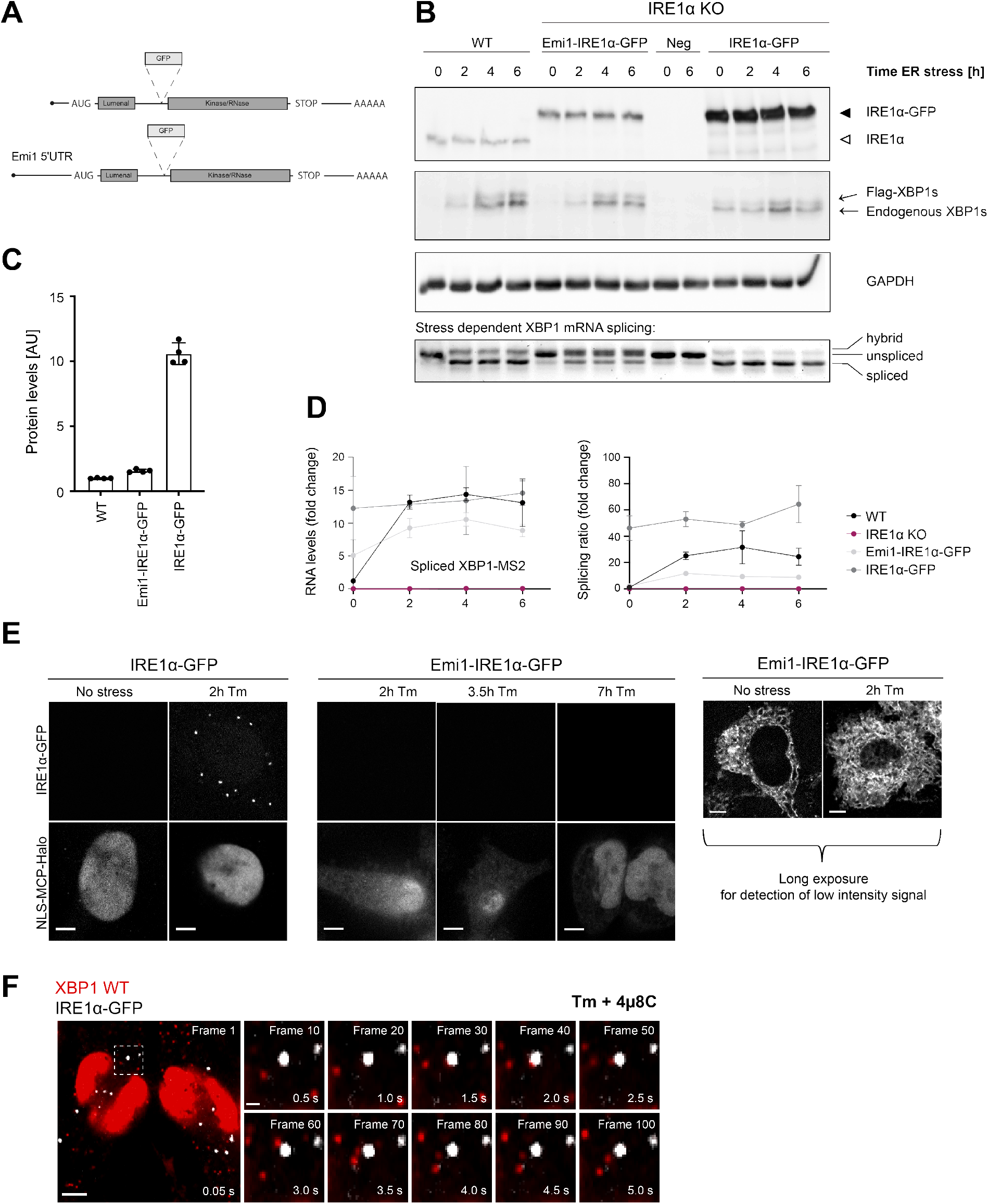
IRE1α is able to splice XBP1u mRNA in the absence of foci formation. **(A)** Schematic representation of IRE1α-GFP construct design analogous to **(Belyy et al., 2019)**. To reduce expression of IRE1α-GFP to match endogenous levels, part of the Emi1 5’ UTR was inserted upstream of the IRE1α-GFP ORF. **(B)** HeLa cells (WT or IRE1α knock-out) expressing either no IRE1α (neg) or reconstituted IRE1α-GFP at low levels (Emi1-IRE1α-GFP) or at high levels (IRE1α-GFP) were kept untreated or treated with 100 nM TG for indicated time points. Upper panels: Western Blot analysis of IRE1α and XBP1s levels in response to TG treatment. XBP1s immunodetection identifies two bands, a lower one corresponding to endogenous XBP1s and an upper one corresponding to the murine, FLAG-tagged XBP1s reporter protein. GAPDH (run in a different gel) was used as a loading control. Bottom panel: Semiquantitative analysis of splicing of WT XBP1 mRNA. Total RNA was isolated from cells that were treated with TG as described above and subjected to RT-PCR with primers flanking the XBP1 intron. Lower band = spliced XBP1, middle band = unspliced XBP1, upper band = hybrid splicing intermediate (one strand spliced, one strand unspliced) **(C)** Quantification of the IRE1α expression levels in cell lines shown in (B). Revert staining of Western blot membranes was used as a normalization value. **(D)** Quantitative RT-PCR to determine the levels of spliced XBP1 mRNA and splicing ratios for the same RNA samples as shown in (B). **(E)** Representative live-cell images of the HeLa cell lines introduced in (C). In cells overexpressing IRE1α-GFP, foci can already be detected at 2h treatment with 5 μg/ml TM. But there are no detectable IRE1α-GFP foci even after prolonged exposure to 5 μg/ml TM under standard imaging conditions in the Emi1-IRE1α-GFP cells. Only long exposure times allow detection of low intensity GFP signal outlining the ER in the absence and presence of 5 μg/ml tunicamycin. **(F)** Representative live-cell images of XBP1 WT reporters (red) in HeLa cells expressing NLS-stdMCP-stdHalo and IRE1α-GFP (gray) under ER stress (5 μg/ml TM) and IRE1α inhibition (50 μM 4μ8C). Dashed box indicates inset that is magnified and shows individual frames of the image series in the right part of the panel. The time series illustrates how individual mRNA particles (red) come close to IRE1α-GFP foci (gray) but do not associate stably nor accumulate in foci. All scale bars = 5 μm, except in single frame magnifications = 1 μm.

IRE1α has been shown to form large oligomeric assemblies and microscopically visible clusters upon induction of ER stress in a number of studies ((Belyy et al., 2019; Kimata et al., 2007; Li et al., 2010; Tran et al., 2021)). Yet, the physiological relevance of these clusters is still not clear. To determine if cluster formation was an artifact of IRE1α overexpression, we aimed to tune IRE1α-GFP expression to match endogenous levels and took advantage of the previously characterized Emi1 5’UTR that has been shown to down-regulate translation approximately 40-fold (Yan et al., 2016). Since this 5’UTR was derived from the cell cycle protein Emi1, we termed the construct Emi1-IRE1α-GFP. For comparison, we also generated an IRE1α-GFP expression construct that was lacking the Emi1 5’UTR and expressed the IRE1α-GFP at higher levels.

As anticipated, Western blot analysis confirmed that reconstituted IRE1α at low (Emi1-IRE1α-GFP) as well as at high levels (IRE1α-GFP) restored the functionality of IRE1α in KO cells, albeit to different extents (Figure 4B). While Emi1-IRE1α-GFP levels were similar to those of endogenous IRE1α, IRE1α-GFP was approximately 10-fold higher expressed (Figure 4C). In both cell lines, ectopic IRE1α-GFP expression rescued XBP1 mRNA splicing under ER stress conditions, as determined by quantitative RT-PCR (Figure 4D) and by Western blot detection of the resulting XBP1s protein (Figure 4B). In line with previous reports (Li et al., 2010), strong overexpression of IRE1α-GFP triggered XBP1 mRNA splicing (and XBP1s synthesis) even in the absence of ER stress, underscoring the importance of adequate IRE1α expression levels for fine tuning of the UPR.

In order to investigate if IRE1α clusters were the sites of XBP1 mRNA splicing on the ER, we imaged IRE1α-GFP in the HeLa cell lines stably expressing XBP1 reporter transcripts along with NLS-stdMCP-stdHalo and Sec61b-SNAP introduced above (Figure 4E). In agreement with earlier reports (Li et al., 2010), we detected IRE1α-GFP foci in cells expressing high levels of the fusion protein even after relatively short induction of ER stress with TM (5 μg/ml) for 2h. Surprisingly, this was not the case for the lowly expressing Emi1-IRE1α-GFP cells, where we were not able to detect IRE1α-GFP clusters even after prolonged exposure to tunicamycin (5 μg/ml) for up to 7h. To make sure that we were not missing IRE1α clusters due to imaging conditions optimized for detection of fast moving mRNA particles (e.g. short 50 ms exposure times), we acquired IRE1α-GFP signal from the same cells in the presence and absence of ER stress but this time using longer exposures (2000 ms) and maximum laser intensities. Under such conditions, we were able to detect IRE1α-GFP signal, which exhibited a characteristic ER-like distribution pattern, but no IRE1α clusters (Figure 4E, right panel).

This observation was intriguing, since it suggested that IRE1α clusters were not necessary for the production of XBP1s, which we were able to detect in the absence of cluster formation (Figure 4D). We wanted to make sure that we were not missing a potential function of the previously observed IRE1α foci and therefore proceeded to image XBP1 WT mRNA recruitment to these oligomeric assemblies at high temporal and spatial resolution (Figure 4F, Supplementary Movie 8). Interestingly, we did not find XBP1 WT transcripts accumulating in IRE1α-GFP clusters even after prolonged TM treatment (5 μg/ml for up to 4h) and inhibition of IRE1α cleavage activity. XBP1 particles freely diffuse around IRE1α-GFP foci and only very rarely colocalize with the IRE1α-GFP signal (Supplementary Movie 9). This is true for XBP1 WT reporter transcripts (in the presence of 4μ8C) as well as for the unspliceable reporter (Supplementary Movie 10).

Our data indicate that Emi1-IRE1α-GFP supports splicing in the absence of foci formation. This suggests that XBP1 mRNA is spliced by lower oligomeric assemblies of IRE1α molecules, which can easily contact ER-associated ribosome-mRNPs, while IRE1α foci or large oligomeric clusters are not the sites of XBP1 mRNA processing during the UPR.

## Discussion

In this study, we investigated the recruitment of XBP1 mRNA to ER-localized IRE1α, which is a fundamental step of the XBP1 splicing mechanism. Based on the organization of such an encounter in yeast and on the visualization of overexpressed IRE1α, splicing of XBP1 mRNA has been attributed to large clusters of IRE1α oligomers that form during the UPR and could function as ER stress response centers (Li et al., 2010). Unexpectedly, our findings challenge this view and suggest a different model for mammalian cells, where IRE1α could be recruited to the XBP1 mRNA, and not the other way around.

Direct visualization of the recruitment of XBP1 mRNAs to the ER surface using a single-molecule imaging approach revealed that XBP1 molecules become ER-associated in an HR2-dependent manner that is consistent with the targeting model proposed by Kohno et al. (Kanda et al., 2016; Yanagitani et al., 2009, 2011). Furthermore, assessment of the translational status of single mRNA particles demonstrated that their ER association depends on interactions with the ribosome-nascent chain complex. Co-translational membrane tethering therefore immobilizes XBP1 transcripts on the ER surface and hints at a substrate recruitment mechanism where IRE1α diffuses through the ER membrane until it encounters unspliced XBP1 mRNAs at the Sec61 translocon. Direct interactions that have been reported for IRE1α and the translocon, SRP, as well as ribosomal RNAs (Acosta-Alvear et al., 2018; Plumb et al., 2015) further increase the affinity of the interaction and underline the potential significance of such a recruitment mechanism.

Upon induction of ER stress, XBP1 transcripts are spliced and released from the ER surface. However, even though we demonstrate that ER association correlates with IRE1α cleavage activity, we did not find XBP1 mRNAs colocalizing with IRE1α clusters. Moreover, large, microscopy-visible clusters were only detected when IRE1α-GFP was overexpressed at high levels. Thus, IRE1α foci are not the primary sites of XBP1 splicing. Instead our findings support a model where ER-poised XBP1 transcripts are processed by functional IRE1α assemblies that are homogenously distributed throughout the ER membrane.

These observations are in good agreement with a parallel study showing that endogenously-tagged IRE1α also fails to assemble into large clusters upon induction of ER stress (Belyy et al., 2021). In this work, the authors characterize IRE1α oligomerization during ER stress and find that the resting pool of IRE1α in the ER membrane is dimeric while in response to stress, transient IRE1α tetramers are assembled as the functional subunits that are required for trans-autophosphorylation and XBP1 splicing. Most likely, such a dynamic equilibrium between dimers and small oligomers allows cells to build timely, finely-tuned responses to local or transient perturbations in ER protein folding.

In combination, our findings suggest a novel mechanism for XBP1 recruitment to functional IRE1α assemblies that are homogenously distributed throughout the ER membrane. Under such circumstances, the enzyme can continuously patrol the membrane to encounter substrates that are targeted there.

Following a different strategy, yeast Ire1p foci arrange the recruitment of unspliced *HAC1* mRNA to the ER membrane and efficiently localize the mRNA for splicing (Aragón et al., 2009; van Anken et al., 2014). Given the strong conservation of most UPR principles, upon visualization of large IRE1α clusters in human cells (Li et al., 2010; Tran et al., 2021) it was plausible to speculate that polarization of IRE1α might build splicing centers. However, these studies mostly relied on overexpression of ectopic IRE1α, which likely contributed to the perception that clusters were required for XBP1 splicing and explains the discrepancies between this and previous reports.

Our findings shed light on a fundamental step of the XBP1 splicing mechanism, which is the recruitment of IRE1α to ER-localized XBP1 transcripts. Yet, several open questions remain:

1. Are XBP1 mRNAs that are tethered to the ER surface as part of ribosome-nascent chain complexes continuously translated? Or is translation stably stalled while the mRNA remains poised for recruitment by IRE1α? And if so, how is translation resumed? And how relevant is the translational status of XBP1 mRNA for splicing?
2. Does IRE1α also bind XBP1 transcripts in the absence of the ribosome/translocon interaction? Since we observe a low degree of splicing for HR2-mutant reporters, we speculate that translation-independent recruitment of XBP1 transcripts and recognition through IRE1α might also be possible.
3. How does IRE1α discriminate between its distinct substrates? Beyond XBP1 splicing, IRE1α processes a broad range of substrates including RIDD (Regulated IRE1-Dependent Decay) mRNAs (Hollien et al., 2009; Hollien & Weissman, 2006) and a recently described, larger group of mRNAs that are processed through an unanticipated mode of cleavage with looser specificity (RIDDLE) (Thomas et al., 2021). Most of these mRNA substrates encode signal sequence-containing proteins and are delivered to the ER by SRP. While we know that activation of the IRE1α RNAse domain requires IRE1α dimerization/oligomerization as well as transautophosphorylation, the specific role and substrate specificity of the distinct assemblies remains unclear. It is tempting to speculate that the interplay of different oligomeric IRE1α assemblies with the translocon/ribosome/SRP environment may define the code for selective processing of distinct IRE1α substrates, avoiding the detrimental cleavage that might result from unrestrained RNA degradation.

In summary, our data have allowed us to visualize and uncover unanticipated features of one of the key steps of UPR initiation, the encounter of XBP1 mRNA with IRE1α to undergo splicing. Additional studies will be needed to further dissect the underlying mechanisms behind the regulation of IRE1α activity in homeostasis and disease.

## Materials and Methods

### DNA constructs

The Gaussia luciferase reporter was the same as previously described (Voigt et al., 2017). Using the same plasmid backbone, we generated a XBP1 wild-type reporter (WT), expressing a N-terminally FLAG-tagged *M. musculus* XBP1 coding sequence and 3’ untranslated region (3’ UTR covers nucleotides 1-948, considering +1 as the first nucleotide after the unspliced mRNA stop codon), followed by 24 MS2 stem loops. Nuclear introns were inserted in this construct to facilitate stability, nuclear export and translation of the reporter mRNA.

HR2 Mutant, spliced and unspliceable constructs were generated by site-directed mutagenesis of WT RNA. In the HR2 mutant, one A nucleotide was inserted 45 nucleotides downstream the 3’ splice site of murine XBP1. This insertion facilitates a translational frameshift that prevents HR2 synthesis, such that the amino acid sequence of the unspliced HR2 mutant protein is identical at the C-terminus to wildtype XBP1s. The spliced reporter plasmid is identical to WT but lacking the 26-nucleotide UPR intron. The unspliceable mutant bears point mutations at the 5’ and 3’ splice site loops. Almost invariant through evolution, positions 1, 3 and 6 of the splice site loops follow the consensus CNGNNGN (Gonzalez et al., 1999; Hooks & Griffiths-Jones, 2011). Mutation of either of these nucleotides disrupts IRE1α cleavage *in vitro* and *in vivo*. Mutations in the 5’ and 3’ splice loops were tCGCAGC and CTaCAGC, respectively (mutation in lowercase).

For the translation reporter of unspliced XBP1 mRNA, a 9xGCN4 spaghetti monster (Eichenberger et al., in preparation) was inserted 35 nucleotides downstream the 3’ splice site, such that the spaghetti monster is in frame with the unspliced polypeptide. In this construct, we removed the last XBP1 nuclear intron, because the insertion of repeats in the close vicinity of its 5’ splice site affected nuclear processing of the transcript.

We used KDEL-Turq2 (Addgene #36204) and Sec61b-SNAP as fluorescent ER markers. Sec61b-SNAP was generated from Addgene construct #121159 (GFP-Sec61b) through replacing the GFP with a SNAP moiety. Single-chain antibodies fused to GFP (scAB-GFP, Addgene #104998) were used for imaging translation sites through nascent polypeptide labeling. NLS-stdMCP-stdHalo (Addgene # 104999) was employed for detection of single mRNA particles.

IRE1α-GFP was generated based on the construct design described by Belyy et al. (2019) and integrated into a phage plasmid for lentiviral expression under the control of a UbiC promotor. To reduce expression levels post-transcriptionally, the Emi1 5’UTR (Yan et al., 2016) was added in front of the ORF. All plasmids are available from the Chao and Aragón labs upon request.

### Cell line generation

HeLa cell lines stably expressing XBP1 and Gaussia luciferase reporter constructs were generated and maintained as previously described (Voigt et al., 2017). Briefly, reporter cassettes were stably integrated into parental HeLa 11ht cells that contain a single FLP site and also express the reverse tetracycline-controlled transactivator (rtTA2-M2) for inducible expression (Weidenfeld et al., 2009). Cells were grown at 37°C and 5% CO_2_ in DMEM + 10% FBS + 1% penicillin, streptomycin (Pen/Strep).

IRE1 was knocked out by CRISPR/Cas9 editing of HeLa cells by transient transfection with the pX459v2-910 plasmid as in (Bakunts et al., 2017) kindly provided by Dr. Eelco van Anken.

NLS-stdMCP-stdHalo, scAB-GFP, KDEL-Turq2, Sec61b-SNAP, IRE1α-GFP and Emi1-IRE1α-GFP fusion proteins were stably integrated into the HeLa cell lines described above via lentiviral transduction. All cell lines were sorted to select for appropriate expression levels for single-molecule imaging.

### Western Blots

For protein extraction, HeLa cell monolayers were washed twiced with ice-cold phosphate saline buffer, and then resuspended directly in Laemmli buffer, supplemented with protease (Complete, Roche) and phosphatase inhibitors. Samples were heated at 95°C for 5 min, loaded on polyacrylamide gels (Thermo Fischer Scientific), and then transferred onto nitrocellulose (GE Healthcare). Protein transfer onto nitrocellulose was confirmed by reversible Ponceau or Revert staining. Immunoblot analysis with was performed using standard techniques. All antibodies used in this study are listed in Supplementary Table 1. Loading correction of immunoblot signals was performed by using GAPDH or tubulin signals as controls, or by quantifying Revert fluorescence after transfer.

Detection of immunolabeled proteins was performed using a commercial chemiluminescent assay (ECL prime; Amersham). Visualization and quantitative measurements were made with a CCD camera and software for Western blot image analysis (Odissey Fc Imager System and Image Studio Lite v 4.0, respectively; Li-COR, Bad Homburg, Germany).

### RNA analysis

RNA was extracted from cells by the guanidine isothyocyanate and phenol-chloroform method (TRIzol; Invitrogen). One μg of total RNA was treated with DNAse I and used for subsequent retrotranscription. 50 to 100 ng of total cDNA was used in real time PCR using SybrGreen (BIORAD). Sequence of primers used in real time PCR are detailed in Supplementary Table 2. For semi-quantitative assessment of splicing, primers flanking the XBP1 intron that specifically amplify murine but not endogenous human XBP1 mRNA PCR products were resolved on 3% agarose gels.

### Membrane flotation assay

For flotation assays, we followed the method originally described by Mechler and Vassalli (Mechler & Rabbitts, 1981). Five minutes before harvesting, subconfluent monolayers of cells were treated with 50 μg/ml cycloheximide (Sigma) to prevent ribosomal runoff from mRNAs. Cultures were washed twice with chilled phosphate saline buffer and resuspended in hypotonic buffer medium [10 mM KCl, 1.5 mM MgCl_2_, 10 mM Tris-HCl pH7.4, 50 μg/ml cycloheximide and a protease inhibitor cocktail (Complete, Roche). Cells were allowed to swell for 5 minutes on ice, and then ruptured mechanically with a Dounce tissue grinder and spun for 2 minutes at 1000 x g and 4°C. The supernatant of this centrifugation was mixed with a 2.5 M sucrose, previously dissolved in TKM buffer [50 mM Tris-HCl pH7.4, 150 mM KCl, 5 mM MgCl_2_, 50 μg/ml cycloheximide plus protease inhibitors], reaching a 2.25 M sucrose concentration. This mixture was layered over 1.5 ml of 2.5M-TKM in a SW40 polyallomer ultracentrifugation tube. On top of the extract-sucrose mix, we layered 6 ml of 2.05 M sucrose-TKM and 2.5 ml of 1.25 M sucrose-TMK. After centrifugation for 10 hours at 25000 rpm in a SW40 Ti Beckman rotor, 1.5 ml fractions were collected from the top to the bottom of the tube and subjected to RNA and protein analysis.

### Single-molecule fluorescence in situ hybridization and immunofluorescence (smFISH-IF)

High precision glass coverslips (170 μm, 18 mm diameter, Paul Marienfeld GmbH) were placed in a 12-well tissue culture plate, HeLa cells were directly seeded on top at 0.5 x 10^5^ cells per well and grown for 48 h. Reporter expression was induced by addition of doxycycline (dox) to the medium for 2h. To ensure strong ER association phenotypes, dox was removed from the medium after that and cells were grown for another 2h until fixation. For ER stress conditions, 5 μg/ml tunicamycin (TM) was added at induction and remained in the medium until fixation.

Combined smFISH-IF was performed as described previously (Dave et al., 2021). Briefly, single-molecule RNA detection was done using Stellaris FISH probes labeled with Quasar 570 (Biosearch Technologies) and designed against the 5’ end of the *M.musculus* XBP1 ORF (Supplementary Table 3). HeLa cells were fixed with 4% paraformaldehyde (Electron Microscopy Sciences) for 10 minutes and then permeabilized in 0.5% Triton-X for another 10 minutes. Cells were pre-blocked in wash buffer (2xSSC (Invitrogen), 10% v/v formamide (Ambion), 3% BSA (Sigma)) for 30 minutes at room temperature, before they were washed twice in 1x PBS and hybridized in hybridization buffer (150 nM smFISH probes, 2xSSC, 10% v/v formamide, 10% w/v dextran sulphate (Sigma)) containing 1:1000 diluted anti-GFP antibody (Aves labs, GFP-1010) for 4 hours at 37°C. After hybridization, cells were washed twice with wash buffer for another 30 minutes each before incubation with anti-chicken IgY secondary antibody conjugated with Alexa-fluor 488 (1:1000 in 1x PBS, Thermo Fisher, A-11039) for 30 minutes. Last, coverslips were washed twice in 1x PBS and then mounted on microscopy slides using ProLong Gold antifade reagent incl. DAPI (Molecular Probes).

smFISH-IF images were acquired on an inverted Zeiss AxioObserver7 microscope that is equipped with a Yokogawa CSU W1 scan head, a Plan-APOCHROMAT 100x 1.4 NA oil objective, a sCMOS camera and an X-Cite 120 EXFO metal halide light source. Z-stacks were acquired in 0.2 μm steps. The exposure times were 500 ms for Quasar 570 and 100 ms for the DAPI channel at maximum laser intensities while the IF signal was acquired at 20 % 488 nm laser intensity for 200 ms.

### smFISH-IF data analysis

Detection of single mRNA as well as translation site spots from fixed cell imaging experiments was performed in KNIME (Berthold et al., 2009) as described before (Voigt, Eglinger, et al., 2019; Voigt, Gerbracht, et al., 2019).

Briefly, individual slices were projected in maximum intensity projections. mRNA and translation site spots were then separately detected using a custom-built KNIME node that runs the spot detection module of TrackMate (Tinevez et al., 2016) in batch mode. This node is available in the KNIME Node Repository: KNIME Image Processing / ImageJ2 / FMI / Spot Detection (Subpixel localization). Detected mRNA and translation site spots in each channel were then co-localized using a nearest neighbor search to link mutual nearest neighbors between the two channels using a distance cut-off of 3 pixels. Nuclear segmentation was performed on the DAPI signal using the Otsu thresholding method while cytoplasmic segmentation was done using the smFISH background signal in the Q570 channel and a manual intensity threshold.

### Live-Cell Imaging

For live-cell imaging, cells were seeded in 35 mm glass-bottom μ-Dishes (ibidi GmbH) 48h prior to the experiment. Depending on the type of experiment, SNAP and Halo fusion proteins were labeled with JF549 or JF646 dyes (HHMI Janelia Research Campus) (Grimm et al., 2015) or SNAP-Cell Oregon Green^®^ (NEB, S9104S).

XBP1 mRNA expression was induced by addition of 1 μg/ml doxycycline to the medium. After 1–2 h, doxycycline was removed to allow proper localization of XBP1 mRNAs to the ER membrane. To inhibit translation, cells were treated with 100 μg/mL puromycin that was added to the cells directly prior to imaging. To induce ER stress, cells were treated with 5 μg/ml tunicamycin (TM) that was added together with doxycycline at induction of mRNA expression and remained in the imaging medium throughout the entire experiment. Similar as for TM, but to inhibit IRE1α activity, the small molecule inhibitor 4μ8C was added at 50 μM together with doxycycline and remained in the medium throughout the imaging experiment. Image acquisition was started not earlier than 1-2 h after doxycycline wash-out to allow proper localization of mRNA molecules and/or induction of the UPR.

Samples were imaged on an inverted Ti2-E Eclipse (Nikon) microscope equipped for live cell imaging and featuring a CSU-W1 scan head (Yokogawa), 2 back-illuminated EMCCD cameras iXon-Ultra-888 (Andor) and an MS-2000 motorized stage (Applied Scientific Instrumentation). Illumination was achieved through 561 Cobolt Jive (Cobolt), 488 iBeam Smart, 639 iBeam Smart (Toptica Photonics) lasers and a VS-Homogenizer (Visitron Systems GmbH). We used a CFI Plan Apochromat Lambda 100X Oil/1.45 objective (Nikon) that resulted in a pixel size of 0.134 μm. For all dualcolor experiments, cells were imaged in both channels (single particles and ER) simultaneously and acquiring fast image series (20 Hz, 100 frames) in a single plane with two precisely aligned cameras. To correct for camera misalignment and chromatic aberrations, images of fluorescent TetraSpeck™ beads were acquired at each imaging session. Cells were maintained at 37°C and 5 % CO_2_ throughout the entire experiment.

### Correlated Diffusion and ER Co-localization Analysis

Images of TetraSpeck™ beads were used to quantify the channel shift in affine transformation mode using the Descriptor-based registration plugin (Preibisch et al., 2010) in Fiji (Schindelin et al., 2012). The transformation model obtained after aligning of the bead images was then re-applied to translate the single mRNA/translation site channel onto the ER channel using as custom-made Fiji macro (Mateju et al., 2020).

Single-particle diffusion and ER co-localization analysis was performed as described before (Voigt et al., 2017). Briefly, we used the KNIME Analytics Platform (Berthold et al., 2009) and a data processing workflow that is available from the Chao lab upon request. The KNIME data analysis pipeline allows segmentation of the ER signal through trainable pixel classification using ilastik (S. Berg et al., 2019). The resulting probability maps are transformed to binary images, which are in turn used to generate distance maps that attribute intensity values to each pixel position with respect to its distance to the closest ER boundary. Positions on the ER are given positive values, while positions away from the ER are defined as negative values. The workflow further correlates mRNA positions (X and Y coordinates) obtained from single particle tracking (SPT) to the ER boundaries at any time point throughout the experiment and computes a cumulative ER localization index through addition of all intensity values that correspond to the positions assumed by a transcript over the experimental time course. To obtain a measure for particle mobility, the workflow further determines instantaneous diffusion coefficients (IDCs) for each track. They are calculated as the mean of all displacements measured by SPT over 100 frames (H. C. Berg, 1993) and can be computed by a custom-made component node that is also available from the KNIME hub (KNIME Hub > Users > imagejan > Public > fmi-basel > components > Instantaneous Diffusion Coefficient).

ER association was quantified for all particles that could be tracked for at least 30 frames and was performed based on IDCs and cumulative ER colocalization indices as described before (Voigt et al., 2017). Values were plotted as scatter plots using the ggplot2 package in R. For the quantification of the degree of ER association per cell, only cells including at least three tracks were included. The analysis was also performed in KNIME and box plots were generated using the ggplot2 and ggpubr packages in R. Data overview and statistics for all live imaging experiments is summarized in Supplementary Table 4.

## Acknowledgements

This work was funded by the Ministerio de Ciencia e Innovación (PID2020-120497RB-I00/ financiado por MCIN/ AEI /10.13039/501100011033) (T.A.), the Novartis Research Foundation (J.A.C.) and a Boehringer Ingelheim Fonds PhD fellowship (T.H.). The authors thank L. Gelman, L.Plantard and J.Eglinger (FMI) for microscopy and image analysis support and H. Kohler (FMI) for cell sorting. We thank Urs Greber and Maite Huarte for critical reading of the manuscript and all members of the Chao and Aragón labs for their input and support.

## Author contributions

T.A. and F.V. conceived the project and designed experiments. S.G.P., R.F. and I.Z. performed molecular biology experiments. T.H. performed smFISH experiments. S.G.P. and R.F. performed flotation assays. F.V. performed live imaging experiments and analyzed data. T.A. and F.V. wrote the manuscript with input from all other authors.

## Supplementary Figures

**Supplementary Figure 1.**
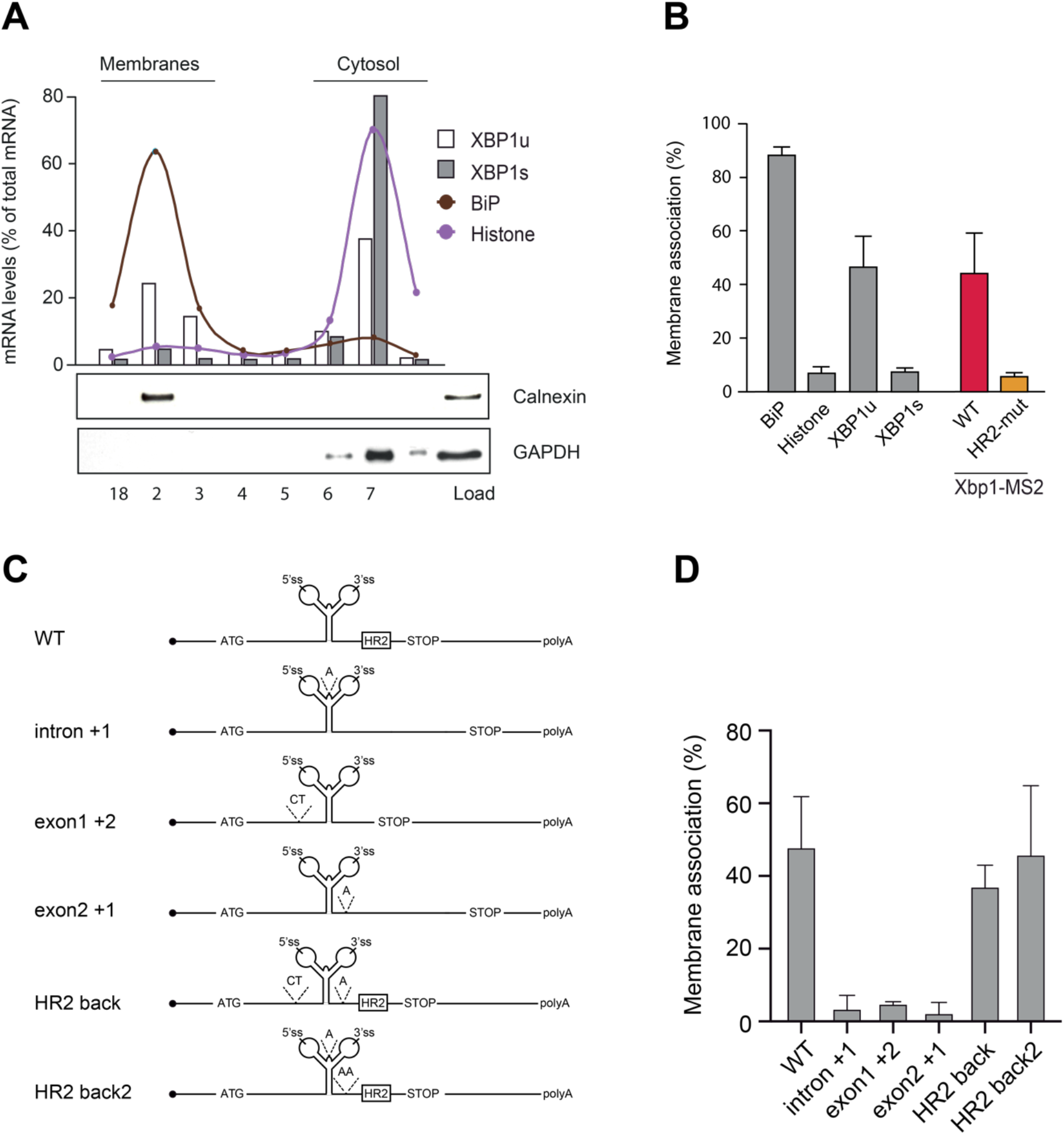
Flotation assays to investigate HR2-mediated recruitment of XBP1 reporter transcripts to ER membranes. **(A)** HEK293 cells were subjected to hypotonic lysis and cytosolic extracts were subjected to flotation in discontinuous sucrose gradients. Fractions 2-3 include floating membranes (as indicated by the ER protein Calnexin), while cytosolic components concentrate in fractions 6-8 (as indicated by GAPDH). Endogenous XBP1s mRNA predominantly associates with cytosolic fractions, while a significant fraction of XBP1u mRNA is found in membrane fractions, albeit to a lesser extent than SRP-targeted mRNAs, such as BiP. **(B)** Quantification of membrane association for endogenous transcripts and reporter mRNAs. The XBP1 WT reporter (red) behaves similar to the endogenous XBP1u transcript. The XBP1 HR2 mutant reporter (yellow) is not found in the membrane fractions and behaves like the endogenous XBP1s mRNA. **(C)** Cartoon illustration of reporter transcripts used in flotation assays. Constructs “intron+1”, “exon1+2”, and “exon2+1” all introduce frameshift mutations abolishing HR2 expression. Constructs “HR2 back” and “HR2 back2” restore the frameshift introduced above and thereby reconstitute membrane association. **(D)** Reporter constructs depicted in C were used to generate HEK293 Flp-In stable cell lines. Cytosolic extracts from these cultures were separated in flotation experiments and analyzed as shown in (A,B). Bar plot represents the average **±** SEM of at least 3 independent experiments.

**Supplementary Fgigure 2.**
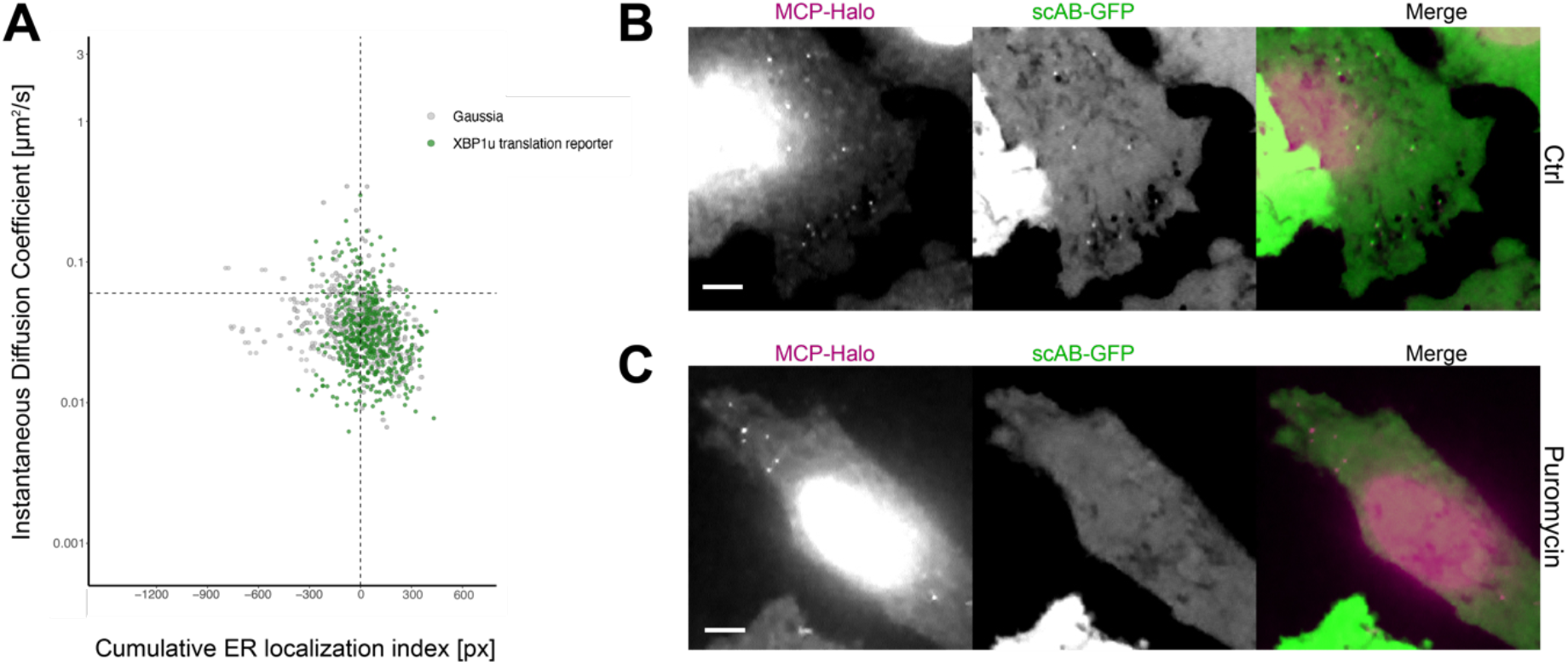
Live imaging of XBP1u translation sites. **(A)** Correlated diffusion and ER colocalization analysis for individual XBP1u translation site tracks (green) and Gaussia mRNA transcripts (gray). Dots are single particles that were tracked for at least 30 frames. Y axis: Instantaneous diffusion coefficients. X axis: Cumulative ER localization index. Positive values indicate ER colocalization. XBP1u translation site tracks scatter similar to Gaussia mRNA tracks. **(B)** Representative live-cell image of XBP1u translation reporter mRNA (magenta) and translation site (green) spots in a HeLa cell stably expressing NLS-stdMCP-stdHalo and scAB-GFP. In the absence of stress, the majority of mRNA transcripts are translated. **(C)** Same as (B) but acquired upon addition of puromycin (100 μg/ml). Translation site spots (green) vanish upon puromycin-mediated translation inhibition. All scale bars = 5 μm.

**Supplementary Figure 3.**
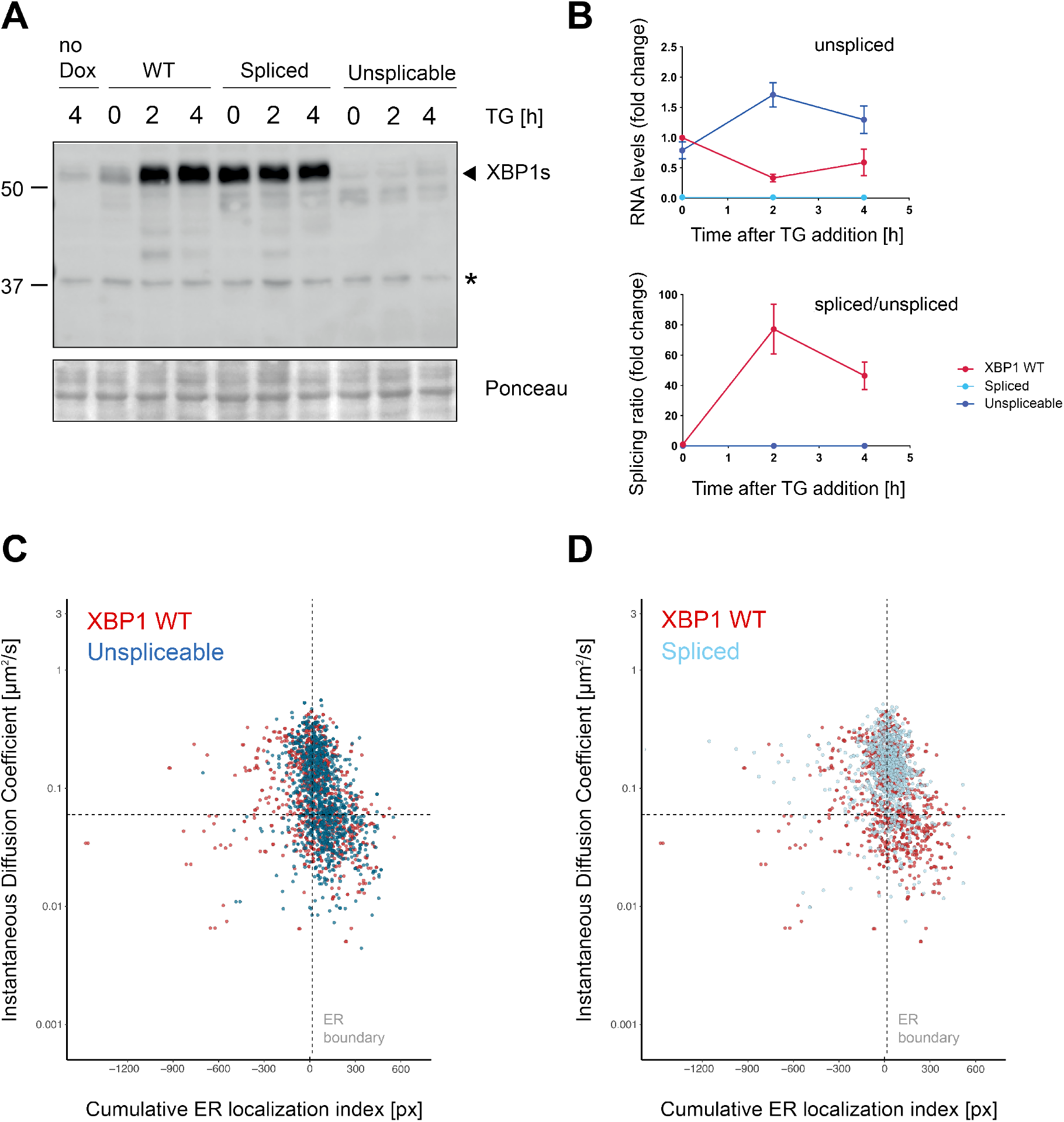
Validation of splice site mutants. **(A)** Western blot analysis of production of spliced XBP1s protein in response to induction of ER stress with 100 nM TG as introduced in Figure 1. Only the WT reporter expressing cells, show a response in XBP1s protein production. Cells expressing the spliced mRNA reporter constitutively produce the 55 kDa XBP1s band, and cells expressing the unspliceable XBP1 reporter fail to produce XBP1s protein at all. **(B)** qPCR-based splicing assays showing the lack of splicing observed for mutant mRNA transcripts (blue) compared to XBP1 WT reporter mRNA (red). **(C)** Correlated diffusion and ER colocalization analysis for XBP1 splice site mutants. Unspliceable reporter transcript (dark blue) compared to XBP1 WT (red). Dots are single particles that were tracked for at least 30 frames. Y axis: Instantaneous diffusion coefficients. X axis: Cumulative ER localization index. Positive values indicate ER colocalization. **(D)** Same analysis as in (A) but for spliced reporter transcripts (XBP1s).

## Supplementary Tables

**Supplementary Table 1.** List of antibodies used for Western blotting

**Supplementary Table 2.** List of primers used for RT-PCR analysis

**Supplementary Table 3.** List of smFISH probes

**Supplementary Table 4.** Imaging data statistics

## Supplementary Movies

**Supplementary Movie 1. XBP1 WT mRNA colocalization with the ER.** HeLa cell line stably expressing XBP1 WT reporter transcripts, NLS-stdMCP-stdHalo and an ER-marker. Simultaneous image acquisition for both channels (XBP1 WT, red, and ER, gray) using 50ms exposure times (100 frames total). The movie is played at 20 fps. The scale bar is 5 μm.

**Supplementary Movie 2. Lack of colocalization with the ER exhibited by XBP1 HR2 mutant transcripts.** HeLa cell line stably expressing XBP1 HR2 mutant reporter transcripts, NLS-stdMCP-stdHalo and an ER-marker. Simultaneous image acquisition for both channels (HR2 mutant, yellow and ER, gray) using 50ms exposure times (100 frames total). The movie is played at 20 fps. The scale bar is 5 μm.

**Supplementary Movie 3. Live imaging of XBP1u translation on the ER.** HeLa cell line stably expressing XBP1u translation reporter transcripts, scAB-GFP and Sec61b-SNAP as ER-marker. Simultaneous image acquisition for both channels (XBP1u translation sites, green, and ER, gray) using 50ms exposure times (100 frames total). The movie is played at 20 fps. The scale bar is 5 μm.

**Supplementary Movie 4. Colocalization of XBP1 unspliceable mutant reporter transcripts with the ER.** HeLa cell line stably expressing XBP1 splice site mutant transcripts, NLS-stdMCP-stdHalo and an ER-marker. Simultaneous image acquisition for both channels (XBP1 Unspliceable, blue and ER, gray) using 50ms exposure times (100 frames total). The movie is played at 20 fps. The scale bar is 5 μm.

**Supplementary Movie 5. Lack of colocalization with the ER exhibited by XBP1 spliced reporter transcripts.** HeLa cell line stably expressing spliced XBP1 transcripts, NLS-stdMCP-stdHalo and an ER-marker. Simultaneous image acquisition for both channels (XBP1 Spliced, light blue and ER, gray) using 50ms exposure times (100 frames total). The movie is played at 20 fps. The scale bar is 5 μm.

**Supplementary Movie 6. Lack of colocalization with the ER exhibited by XBP1 WT transcripts in response to ER stress.** HeLa cell line stably expressing XBP1 WT reporter transcripts, NLS-stdMCP-stdHalo and an ER-marker. Cells were treated with 5 μg/ml tunicamycin for 3-4h prior to image acquisition. Simultaneous image acquisition for both channels (XBP1 WT, red, and ER, gray) using 50ms exposure times (100 frames total). The movie is played at 20 fps. The scale bar is 5 μm.

**Supplementary Movie 7. XBP1 WT mRNA colocalization with the ER during ER stress and inhibition of IRE1α RNase activity.** HeLa cell line stably expressing XBP1 WT reporter transcripts, NLS-stdMCP-stdHalo and an ER-marker. Cells were treated with 5 μg/ml tunicamycin and 50 μM 4μ8C for 3-4h prior to image acquisition. Simultaneous image acquisition for both channels (XBP1 WT, red, and ER, gray) using 50ms exposure times (100 frames total). The movie is played at 20 fps. The scale bar is 5 μm.

**Supplementary Movie 8. No accumulation of XBP1 WT transcripts in IRE1α-GFP foci during IRE1α inhibition.** HeLa cell line stably expressing XBP1 WT reporter transcripts, NLS-stdMCP-stdHalo and IRE1α-GFP. Cells were treated with 5 μg/ml tunicamycin and 50 μM 4μ8C for 2-3h prior to image acquisition. Simultaneous image acquisition for both channels (XBP1 WT, red, and IRE1α-GFP, gray) using 50ms exposure times (100 frames total). The movie is played at 20 fps. The scale bar is 5 μm.

**Supplementary Movie 9. Detection of single XBP1 WT transcripts in IRE1α-GFP foci is possible but extremely rare.** HeLa cell line stably expressing XBP1 WT reporter transcripts, NLS-stdMCP-stdHalo and IRE1α-GFP. Cells were treated with 5 μg/ml tunicamycin and 50 μM 4μ8C for 2-3h prior to image acquisition. Simultaneous image acquisition for both channels (XBP1 WT, red, and IRE1α-GFP, gray) using 50ms exposure times (100 frames total). The movie is played at 20 fps. The scale bar is 5 μm. White arrow indicates a single XBP1 mRNA particle that colocalizes with an IRE1α cluster.

**Supplementary Movie 10. No accumulation of XBP1 splice site mutant transcripts in IRE1α-GFP foci.** HeLa cell line stably expressing XBP1 splice site mutant reporter transcripts, NLS-stdMCP-stdHalo and IRE1α-GFP. Cells were treated with 5 μg/ml tunicamycin for 3-4h prior to image acquisition. Simultaneous image acquisition for both channels (Unspliceable XBP1 reporter, blue, and IRE1α-GFP, gray) using 50ms exposure times (100 frames total). The movie is played at 20 fps. The scale bar is 5 μm.

